# treespat:an R package for spatial tree diversity analysis

**DOI:** 10.1101/2023.05.22.541683

**Authors:** F. Chianucci, N. Puletti, L. Cesaretti

## Abstract

Forest structure is a key element in understanding functionality and resilience of forest ecosystems. Analyses of forest structure and diversity have traditionally employed non-spatial measures, due to the simplicity of such approach. However, spatial structure can provide a deeper understanding of tree patterns, but widespread use of a spatially-explicit approach has been often limited by the complexity of theoretical equations and formulas, and the lack of freely distributed tools to calculate such spatial indices. To fill this gap we created the R package `treespat`, which allows to calculate some of the most diffuse stand-level spatial diversity indices. The document describes theory, basic definition and example application for using the `treespat` package.

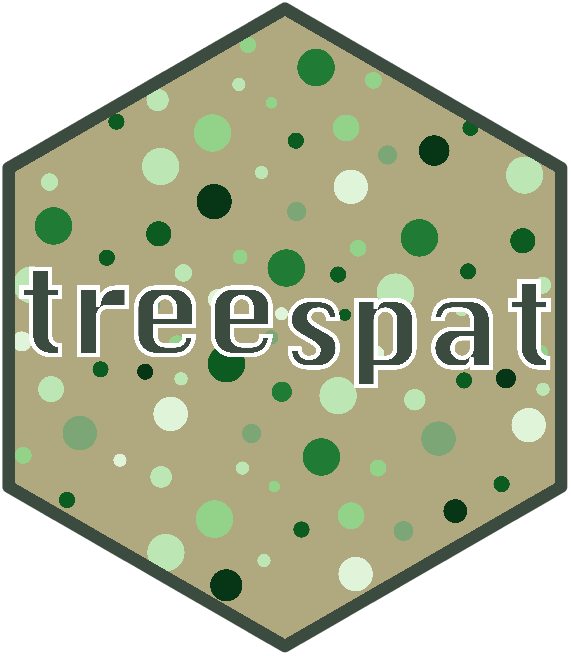

## Introduction

Forest structure is a key element in understanding functionality and resilience of forest ecosystems (Seidl et al., 2014) and a basic element of species diversity (Staudhammer and LeMay, 2001). An increasing complexity of stand structure is often linked to a higher number of species and, consequently, to greater ecological stability.

Analyses of forest structure and diversity have traditionally employed non-spatial measures, due to the simplicity of such approach. However, spatial structure can provide a deeper understanding of tree patterns (Grotti et al. 2019) and determines the properties of the ecosystem, including total biomass production, biodiversity, ecosystem functioning (Gadow et al., 2012; Castaño-Santamaría et al. 2021). However, use of a spatial approach in studying tree diversity is often hampered by the difficulty of implementing theoretical equations and formulas, and the lack of freely distributed tools to calculate such spatial indices.

According to Pommerening (2002), the majority of indices quantify spatial forest structure can be divided into three groups:

1. **stand-level attributes** based on neighborhood relations;
2. **distance-dependent measures** describing stand structure;
3. **continuous functions**.

While some existing R package already implemented some distance-dependent measures (2) and continuous functions (3) as for example in the `spatstat` family packages (Baddeley et al. 2015) and `spatdiv` package (see https://ericmarcon.github.io/SpatDiv), no available R packages allow to calculate stand-level parameters (1). Nonetheless, stand-level spatial indices are important to explore small-scale differences in diversity and also to link stand structure to ecosystem management, functioning and diversity (see example in Grotti et al. 2019).

The R package `treespat` allows to calculate some of the most diffuse stand-level spatial diversity indices, according to work by Pommerening (2002). The next section will describe the basic theory of most diffuse stand-level indices implemented in the package.

### Spatial diversity indices

The package implements three of the most diffuse stand-level spatial indices (Pommerening 2002, Pommerening 2020): *size differentiation, size dominance, species mingling*.

Their implementation requires mapped tree data, namely horizontal tree (x, y) position and an additional mark (*m*) attribute, e.g. species, tree height, tree diameter measured for all individual in a forest-inventoried plot.

Two additional indices were also calculated by the package:

- *Mean Directional Index* is an angle-based index (Corral-Rivas, 2006) representing the spatial arrangement of trees.
- *Stand Complexity Index* is a stand level measure of structural 3D complexity.

Formula of the indices is reported below.

### Size differentiation

An early measure of spatial size inequality in plants was introduced by Gadow (1993) as the mean of the ratio of smaller and larger plant sizes *m* of the *k* nearest neighbors:

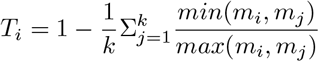

where *m* can be any quantifiable plant size measure, e.g. height or tree diameter among others. The value of *T*_*i*_ increases with increasing average size difference between neighboring trees, with *T*_*i*_ = 0 implying that neighboring trees have equal size. This index is based on the size-ratio construction principle.

#### 2. Size dominance

(Aguirre et al. 2003) proposed an index using the size-comparison construction principle. This method turns the continuous size variables into a binary problem. The dominance index is defined as the mean fraction of trees among the *k* nearest neighbors of a given individual *j* that are smaller than *i*.

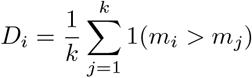

where *m* can be any quantifiable plant size measure, e.g. height or tree diameter among others. The indicator function returns 1, if the size mark of plant *i* exceeds that of neighboring plant *j*, otherwise 0. It can have *k* possible discrete outcomes.

#### 3. Mingling Index

One very intuitive extension of taxonomic species diversity (either richness or abundance) is considering spatial mingling, namely how plants of the same (con-specific neighbors) or different (hetero-specific neighbors) species are arranged in space.

The mingling index (Aguirre et al. 2003) calculates the proportion of the *k* nearest neighbors that do not belong to the same species as the reference tree *i*:

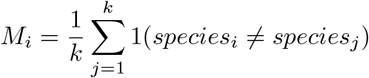

For example, with four neighbors, the mingling attribute *M*_*i*_ can assume five values, ranging from 0 (all trees are of the same species) to 1 (all trees belong to different species).

#### 4. Mean directional index

The mean directional index (Corral-Rivas, 2006) is defined as the sum of the unit vectors from the reference tree *i* to its *k* nearest neighbors and represents the spatial arrangement of tree:

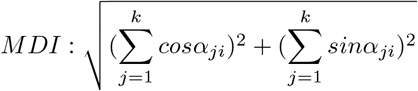

#### 5. Stand Complexity Index (SCI; Zenner and Hibbs, 2000)

The SCI index has been developed to incorporate the 3D structural complexity into spatial diversity analysis. It can be calculated from mapped tree data, as:

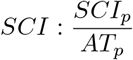

Where *SCI*_*p*_ is the continuous faceted surface, calculated in the 3D-dimensional space, and *AT*_*p*_ is the projected areas of the triangles forming the faceted surface. For the calculation, the 3D-space consider both the horizontal (x, y) tree position and tree height (*z*), but other tree attribute can be considered for the third-dimension.

### Installation

You can install the development version of `treespat` from [GitLab](https://gitlab.com/fchianucci/LAD) using `devtools` (Wickham et al. 2021):

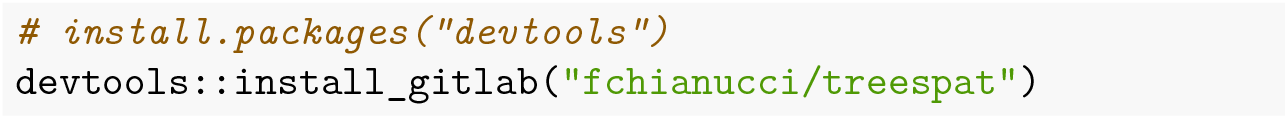

### Basic usage

The package implements the function above by calculating the values for each reference *i* tree and a user-defined number of *k* neighbors (suggested values is *k ≤* 4).

For each index, plot-level estimates were then calculated by calculating their arithmetic mean.

In addition a NN1 edge correction method is also applied for all the indices (with the exception of SCI), following Pommerening and Stoyan (2006).

We will illustrate the calculation of all indices in `treespat` using an example dataset (`bf`) of a semi-natural oak-hornbeam forest sampled in Northern Italy. it provides replicated, repeated measurements at single-tree level conducted in three ∼ 1-ha permanent plots (*Core Areas*, abbreviated as CA); all standing trees in these plots were mapped and inventoried in 1995, 2005, and 2016.

The dataset is described in Fardusi et al. 2018. The original dataset included some coppice stools, which have identical tree coordinates, and dead trees. For usage in `treespat`, data were elaborated to consider only living trees, while for coppice stools we calculated their cumulative basal area, and then we kept only an individual, whose diameter was calculated from inverting the cumulative basal area, and whose height was that of the tallest stool. For details see Puletti et al. (2023).

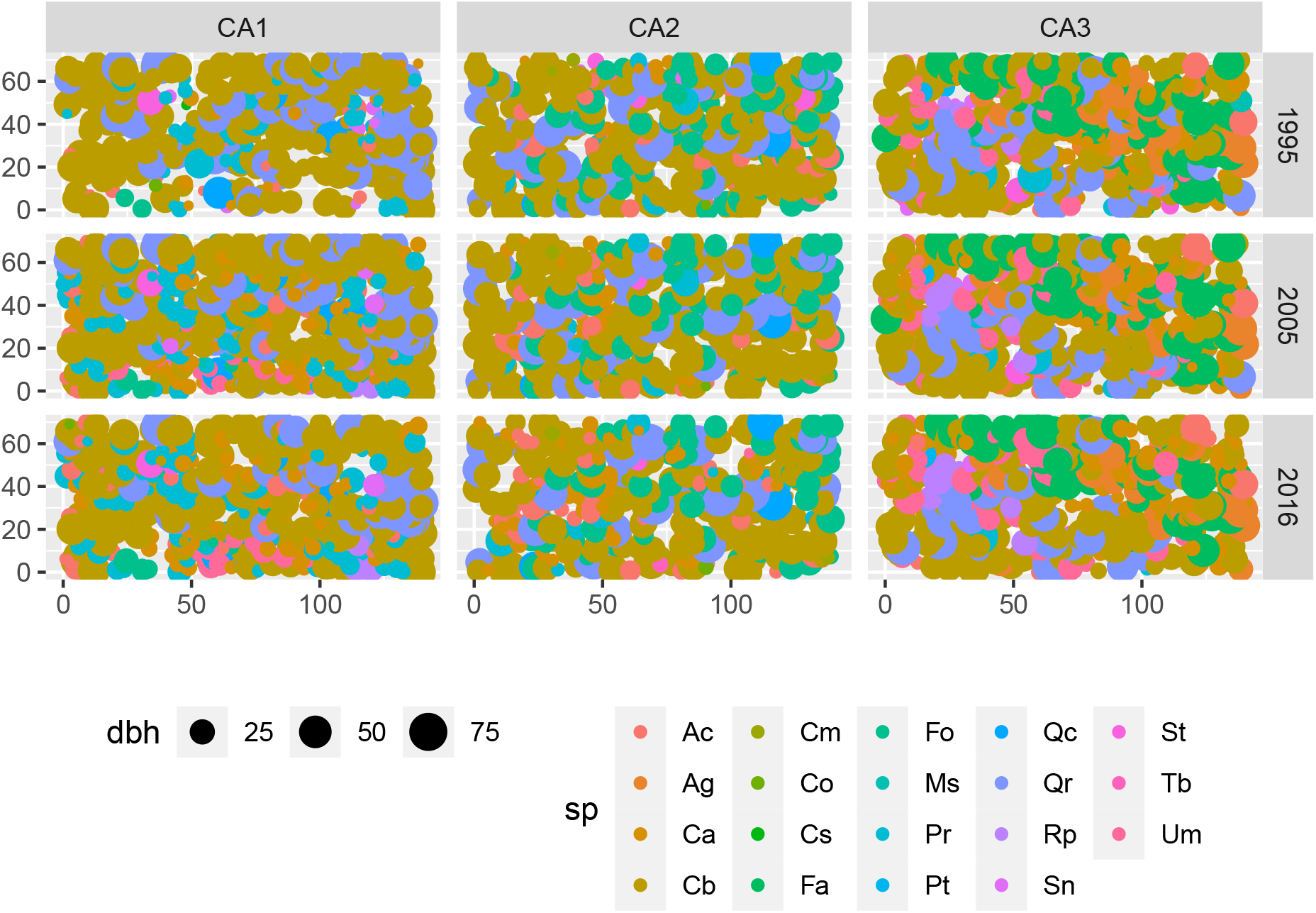

We can inspect the dataset (`bf`):

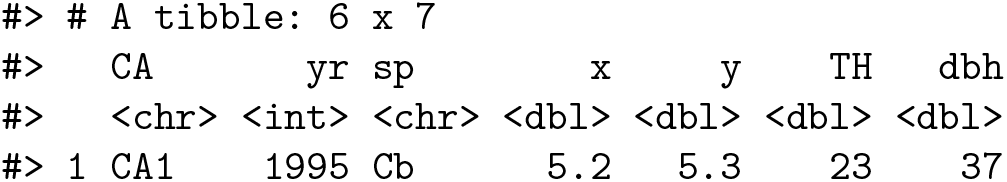
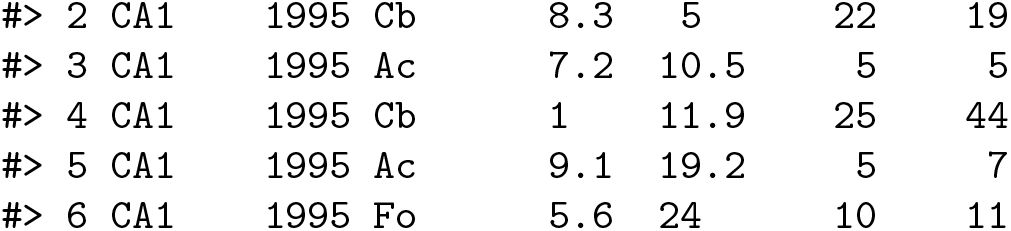

- *CA* is the plot identifier;
- *yr* is the sampling year;
- *sp* is the code representing the tree species;
- *x, y* are the horizontal coordinates from the bottom-left plot origin (0,0), expressed in m;
- *TH* is the tree top height, in m;
- *dbh* is the diameter at the breast height, in cm.

All the spatial functions implemented requires an input dataframe with:

- the tree positions (`.x, .y`);
- a quantitative `.mark` (e.g. diameter or height) or `.species` (for mingling index);
- information about plot size (`xmax`, `ymax` or `radius`) and `shape` (to apply edge corrections);
- the *k* number of neighboring trees to consider (`max.k`);
- a grouping column(s) `.groups`.

Below we provide examples with *k*=3 neighboring trees.

- **Diameter differentiation** (*DIFF*):

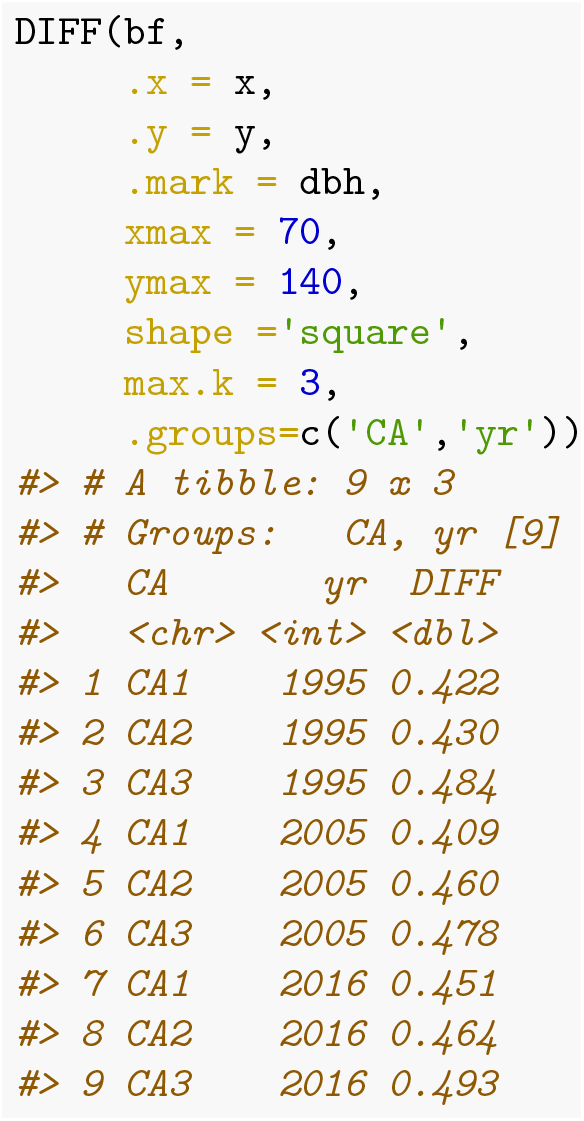

- **Size dominance** (*DDOM*):

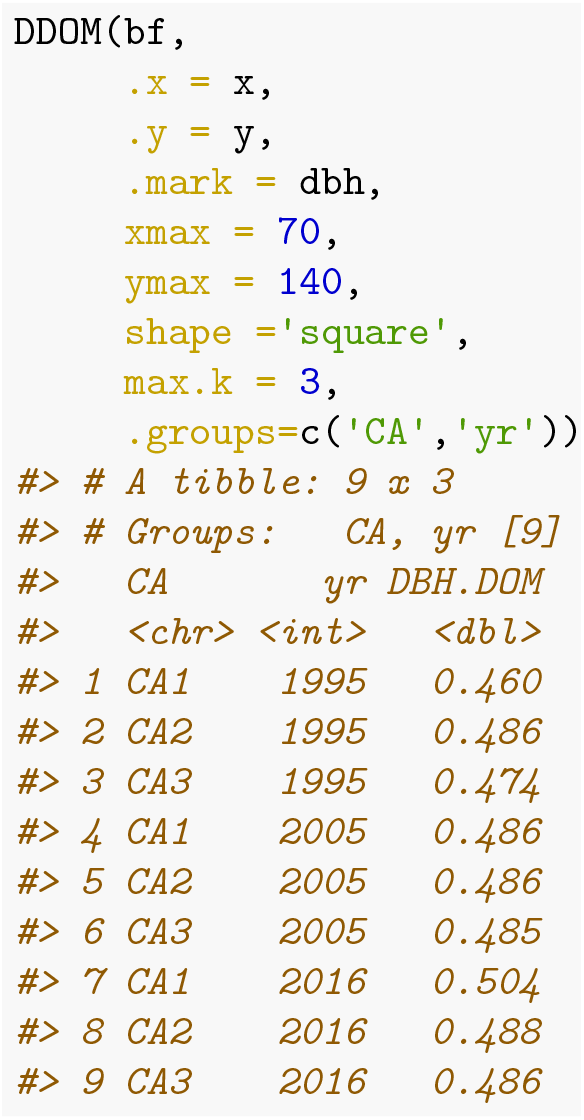

In the above example we used tree diameter, but any other quantitative `.mark` (e.g. tree height) can be used.

- **Mingling index** (*MING*):

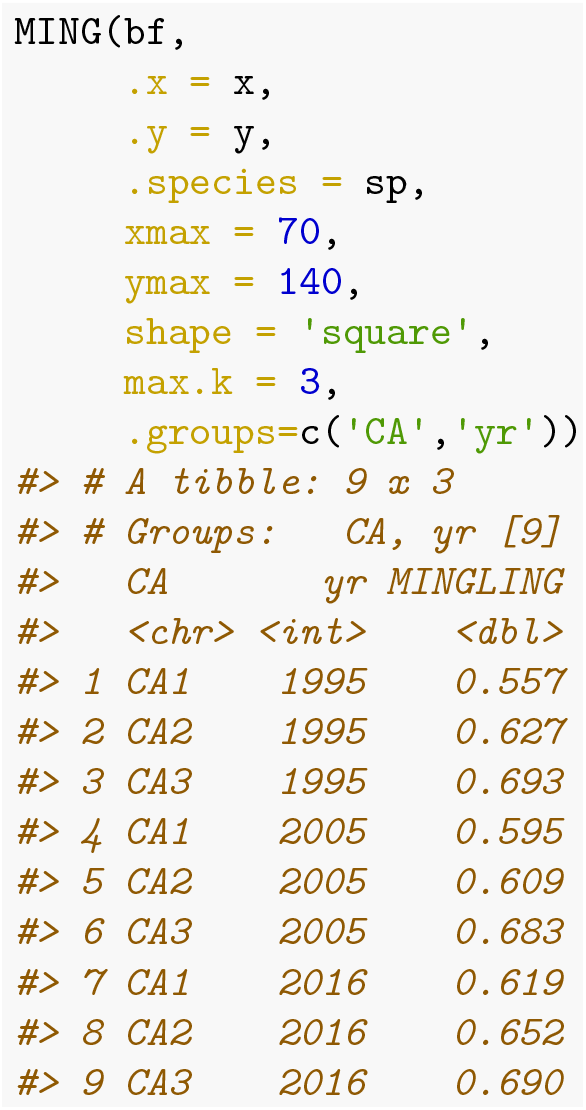

- **Mean directional index** (*MDIR*):

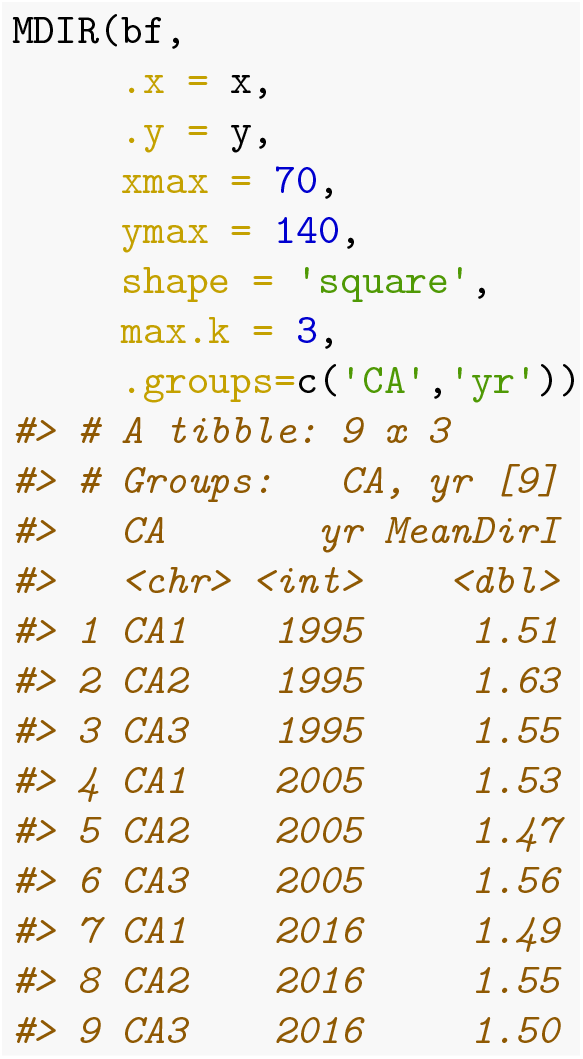

- **Stand complexity index** (*SCI*):

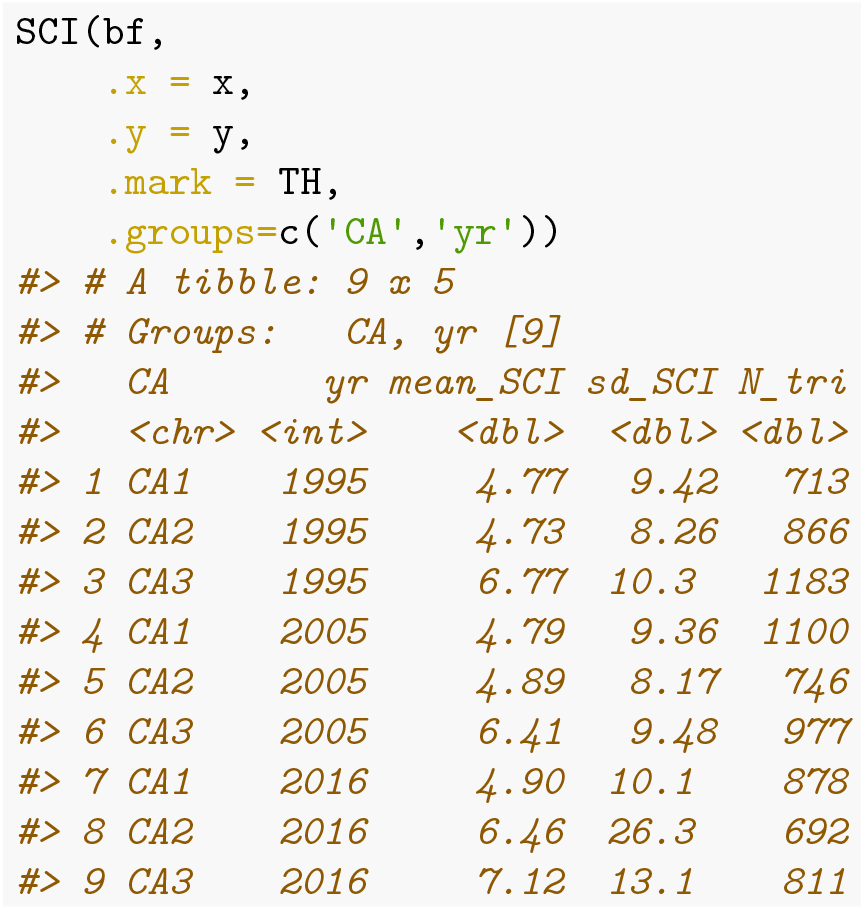

*Note: we did not implemented edge correction in this function as it requires different approach compared to NN1 (see Põldveer et al. 2021)*.

